## Supplementary material for "*Babesia bovis* Rad51 ortholog influences switching of *ves* genes but is not essential for segmental gene conversion in antigenic variation": S1 Table.docx

**S1 Table. Organizational nature of the *ves* loci to which reads mapped.** These mappings are based upon the *Babesia bovis* C9.1 line genome (available at ftp://ftp.sanger.ac.uk/pub/pathogens/Babesia/).

**RNA1: ko1, 1 month**

Region	p-value^1^	Length^2^	Locus type^3^

6915-7268	0.00	354	divergent a/a^4^

782938-783288	0.00	351	unidirectional a

786356-786709	0.00	354	unidirectional a

1011139-1011492	0.00	354	unidirectional a

1039551-1039898	0.00	348	unidirectional a

1055223-1055551	0.00	329	divergent b^4^/a

1122014-1122364	0.00	351	divergent a/a

1208548-1208877	0.00	330	divergent a/b

1399269-1399601	0.00	333	divergent b/a

1403602-1403967	0.00	366	divergent b/a

1404548-1404874	0.00	327	divergent b/a

1600915-1601241	0.00	327	unidirectional a

1618150-1618599	0.00	450	unidirectional a

2678070-2678405	0.00	336	divergent a/a

3137303-3137650	0.00	348	no annotated gene

3396845-3397184	0.00	340	no annotated gene

5260614-5260935	0.00	322	divergent b/a

5456956-5457291	0.00	336	divergent b/a

5458987-5459181	0.00	195	divergent b/a

6205332-6205667	0.00	336	unidirectional a

6218668-6219009	0.00	342	divergent a/a

7443575-7443907	0.00	333	unidirectional a

7762155-7762487	0.00	333	divergent a/b

10 divergent; 8 unidirectional; 2 with no annotated gene (20 total); div/uni= 1.25.

**RNA2: ko1, 5 months**

Region	p-value	Length	Locus type

6915-7268	0.00	354	divergent a/a

285751-286083	0.00	333	divergent a/a

782938-783288	0.00	351	unidirectional a

785282-785626	0.00	345	unidirectional a

786356-786709	0.00	354	unidirectional a

1011139-1011492	0.00	354	unidirectional a

1037935-1038261	0.00	327	unidirectional a

1039551-1039898	0.00	348	unidirectional a

1055223-1055551	0.00	329	divergent b/a

1122014-1122364	0.00	351	divergent a/a

1133046-1133396	0.00	351	divergent b/a

1208548-1208877	0.00	330	divergent a/b

1399269-1399601	0.00	333	divergent b/a

1403602-1403967	0.00	366	divergent b/a

1404548-1404874	0.00	327	divergent b/a

1600915-1601241	0.00	327	unidirectional a

1618150-1618599	0.00	450	unidirectional a

2678070-2678405	0.00	336	divergent ves/ves

2867318-2867671	0.00	354	divergent a/b

2897991-2898213	8.55E-7	223	unidirectional a

3137303-3137650	0.00	348	no annotated gene

3396845-3397184	0.00	340	no annotated gene

5260614-5260991	0.00	378	divergent b/a

5456956-5457291	0.00	336	divergent b/a

5458987-5459181	0.00	195	divergent b/a

6218668-6219009	0.00	342	divergent a/a

7443575-7443907	0.00	333	unidirectional a

7762155-7762487	0.00	333	divergent a/b

13 divergent; 7 unidirectional; 2 no annotated gene (22 total); div/uni= 1.86

**RNA3: ko2, 1 month**

Region	p-value	Length	Locus type

6915-7268	0.00	354	divergent a/a

782938-783288	0.00	351	unidirectional a

786356-786709	0.00	354	unidirectional a

1011139-1011492	0.00	354	unidirectional a

1039551-1039898	0.00	348	unidirectional a

1064683-1065007	0.00	325	divergent a/b

1133046-1133396	0.00	351	divergent b/a

1208548-1208877	0.00	330	divergent a/b

1399269-1399601	0.00	333	divergent b/a

1402278-1402513	5.49E-15	236	divergent b/a

1403602-1403967	0.00	366	divergent b/a

1600915-1601241	0.00	327	unidirectional a

1618150-1618599	0.00	450	unidirectional a

2678070-2678405	0.00	336	divergent ves/ves

2678790-2678964	9.73E-7	175	divergent ves/ves

3137303-3137650	0.00	348	no annotated gene

3396845-3397184	0.00	340	no annotated gene

5260614-5260991	6.61E-7	378	divergent b/a

5457143-5457291	0.00	149	divergent b/a

5458987-5459181	0.00	195	divergent b/a

6218685-6218996	1.92E-7	312	divergent a/a

7443575-7443907	0.00	333	unidirectional a

7762155-7762487	0.00	333	divergent a/b

10 divergent; 6 unidirectional; 2 no annotated gene (20 total); div/uni= 1.67

**RNA4: ko2, 5 months**

Region	p-value	Length	Locus type

6915-7268	3.52E-15	354	divergent a/a

264245-264601	0.00	357	unidirectional a

286923-287037	0.00	115	divergent a/a

782938-783288	0.00	351	unidirectional a

785381-785471	2.17E-8	91	unidirectional a

786356-786709	0.00	354	unidirectional a

1011139-1011492	0.00	354	unidirectional a

1039551-1039898	0.00	348	unidirectional a

1055223-1055551	0.00	329	divergent b/a

1122014-1122364	0.00	351	divergent a/a

1125758-1125858	1.76E-15	101	divergent ves/a

1208548-1208877	0.00	330	divergent a/b

1399269-1399601	0.00	333	divergent b/a

1402277-1402513	4.81E-9	237	divergent b/a

1403602-1403967	0.00	366	divergent b/a

1404663-1404777	7.26E-7	115	divergent b/a

1600915-1601241	0.00	327	unidirectional a

1618150-1618599	0.00	450	unidirectional a

2897979-2898313	0.00	335	unidirectional a

2918405-2918586	0.00	182	divergent b/ves

3137303-3137650	0.00	348	no annotated gene

3396845-3397184	0.00	340	no annotated gene

5260614-5260991	0.00	378	divergent b/a

5457143-5457291	5.16E-14	149	divergent b/a

5458986-5459181	0.00	196	divergent b/a

7421824-7421883	6.72E-8	60	unidirectional a

7443575-7443907	0.00	333	unidirectional a

7762155-7762487	0.00	333	divergent a/b

12 divergent; 9 unidirectional; 2 no annotated gene (23 total); div/uni= 1.33

**RNA5: ko3, 1 month**

Region	p-value	Length	Locus type

6915-7268	0.00	354	divergent a/a

264245-264601	0.00	357	unidirectional a

285770-286071	7.21E-7	302	divergent a/a

286759-287112	1.90E-7	354	divergent a/a

782938-783288	0.00	351	unidirectional a

785282-785626	0.00	345	unidirectional a

786375-786585	9.81E-6	211	unidirectional a

1011139-1011492	0.00	354	unidirectional a

1028822-1029096	0.00	275	divergent b/a

1039551-1039898	0.00	348	unidirectional a

1122014-1122364	0.00	351	divergent a/a

1133046-1133396	0.00	351	divergent b/a

1208548-1208877	0.00	330	divergent a/b

1399269-1399601	0.00	333	divergent b/a

1403602-1403967	0.00	366	divergent b/a

1404548-1404874	0.00	327	divergent b/a

1600915-1601241	7.98E-14	327	unidirectional a

1618168-1618580	2.83E-6	413	unidirectional a

2678070-2678394	6.75E-10	325	divergent ves/ves

2867329-2867671	8.14E-7	343	divergent a/b

2897979-2898332	0.00	354	unidirectional a

3137303-3137650	0.00	348	no annotated gene

3396845-3397186	0.00	342	no annotated gene

5260614-5260935	3.86E-17	322	divergent b/a

5457143-5457291	0.00	149	divergent b/a

5458074-5458335	0.00	262	divergent b/a

5458987-5459181	0.00	195	divergent b/a

6216891-6217207	6.83E-8	317	divergent a/a

6218668-6219009	0.00	342	divergent a/a

7443575-7443907	0.00	333	unidirectional a

7762166-7762487	1.24E-9	322	divergent a/b

16 divergent; 8 unidirectional; 2 no annotated gene (26 total); div/uni= 2.00

**RNA6: ko3, 5 months**

Region	p-value	Length	Locus type

6915-7268	0.00	354	divergent a/a

286759-287112	0.00	354	divergent a/a (a pair)

782938-783288	0.00	351	unidirectional a

785379-785471	2.08E-11	93	unidirectional a

786356-786709	0.00	354	unidirectional a

1011139-1011492	0.00	354	unidirectional a

1039551-1039898	0.00	348	unidirectional a

1055223-1055551	0.00	329	divergent b/a

1122014-1122364	0.00	351	divergent a/a

1125758-1125858	0.00	101	divergent a/a

1133046-1133396	0.00	351	divergent b/a

1208548-1208877	0.00	330	divergent a/b

1399269-1399601	0.00	333	divergent b/a

1402278-1402513	0.00	236	divergent b/a

1403602-1403967	0.00	366	divergent b/a

1404548-1404874	0.00	327	divergent b/a

1600915-1601241	0.00	327	unidirectional a

1618150-1618599	0.00	450	unidirectional a

2001903-2002139	0.00	237	unidirectional a

2676785-2676969	0.00	185	divergent ves/ves

2678070-2678405	1.45E-10	336	divergent a/a

2897979-2898313	5.69E-14	335	unidirectional a

2918405-2918586	0.00	182	divergent b/ves

3137303-3137650	1.60E-9	348	no annotated gene

3396845-3397184	0.00	340	no annotated gene

5260614-5260991	0.00	378	divergent b/a

5458986-5459181	2.06E-16	196	divergent b/a

7421824-7421883	1.57E-12	60	unidirectional a

7443575-7443907	0.00	333	unidirectional a

7762155-7762487	0.00	333	divergent a/b

14 divergent; 9 unidirectional; 2 no annotated gene (25 total); div/uni= 1.56

**RNA7: CE11(B8), 1 month**

Region	p-value	Length	Locus type

6915-7268	0.00	354	divergent a/a

264245-264601	6.37E-9	357	unidirectional a

782938-783288	0.00	351	unidirectional a

785282-785626	0.00	345	unidirectional a

786356-786709	0.00	354	unidirectional a

1011139-1011492	0.00	354	unidirectional a

1028822-1029096	0.00	275	divergent b/a

1037935-1038261	3.73E-18	327	unidirectional a

1039551-1039898	0.00	348	unidirectional a

1055223-1055551	0.00	329	divergent b/a

1064683-1065007	1.33E-15	325	divergent a/b

1122014-1122364	0.00	351	divergent a/a

1133046-1133396	0.00	351	divergent b/a

1208548-1208877	0.00	330	divergent a/b

1399269-1399601	0.00	333	divergent b/a

1402184-1402513	0.00	330	divergent b/a

1403602-1403967	0.00	366	divergent b/a

1404548-1404874	0.00	327	divergent b/a

1411481-1411704	0.00	224	unidirectional a

1600915-1601241	0.00	327	unidirectional a

1618150-1618599	0.00	450	unidirectional a

2001876-2002139	0.00	264	unidirectional a

2676785-2676969	0.00	185	divergent ves/ves

2678070-2678405	0.00	336	divergent a/a

2867318-2867671	0.00	354	divergent a/b

2882658-2882975	4.26E-10	318	unidirectional a

2897979-2898332	0.00	354	unidirectional a

2918405-2918586	0.00	182	divergent b/ves

3137303-3137650	0.00	348	no annotated gene

3396845-3397186	1.17E-16	342	no annotated gene

5260614-5260991	0.00	378	divergent b/a

5456956-5457291	0.00	336	divergent b/a

5458074-5458335	0.00	262	divergent b/a

5458922-5459254	0.00	333	divergent b/a

6205332-6205667	0.00	336	unidirectional a

6218679-6218996	8.82E-10	318	divergent a/a

7380075-7380237	2.37E-7	163	unidirectional a

7421547-7421883	0.00	337	unidirectional a

7443575-7443907	0.00	333	unidirectional a

7762155-7762487	0.00	333	divergent a/b

16 divergent; 15 unidirectional; 2 no annotated gene (33 total); div/uni= 1.07

**RNA8: CE11(B8), 5 months**

Region	p-value	Length	Locus type

6915-7268	0.00	354	divergent a/a

264245-264601	5.02E-7	357	unidirectional a

782938-783288	0.00	351	unidirectional a

785282-785626	0.00	345	unidirectional a

786356-786709	0.00	354	unidirectional a

1011139-1011492	0.00	354	unidirectional a

1028822-1029096	0.00	275	divergent b/a

1037946-1038261	1.39E-7	316	unidirectional a

1039551-1039898	0.00	348	unidirectional a

1055223-1055551	0.00	329	divergent b/a

1064683-1065007	0.00	325	divergent a/b

1122014-1122364	0.00	351	divergent a/a

1133046-1133396	0.00	351	divergent b/a

1208548-1208877	0.00	330	divergent a/b

1399269-1399601	0.00	333	divergent b/a

1402184-1402513	0.00	330	divergent b/a

1403602-1403967	0.00	366	divergent b/a

1404548-1404874	0.00	327	divergent b/a

1411481-1411704	0.00	224	unidirectional a

1600915-1601241	0.00	327	unidirectional a

1618150-1618599	0.00	450	unidirectional a

2001876-2002139	0.00	264	unidirectional a

2676785-2676969	0.00	185	divergent ves/ves

2678070-2678405	0.00	336	divergent a/a

2867318-2867671	0.00	354	divergent a/b

2882658-2882975	7.82E-13	318	unidirectional a

2897979-2898332	0.00	354	unidirectional a

2918405-2918586	0.00	182	divergent b/ves

3137303-3137650	0.00	348	no annotated gene

3396845-3397184	0.00	340	no annotated gene

5260614-5260991	0.00	378	divergent b/a

5456956-5457291	0.00	336	divergent b/a

5458074-5458335	0.00	262	divergent b/a

5458922-5459254	0.00	333	divergent b/a

6205332-6205667	0.00	336	unidirectional a

6218672-6218996	9.92E-7	325	divergent a/a

7380075-7380237	1.15E-8	163	unidirectional a

7421547-7421883	0.00	337	unidirectional a

7443575-7443907	0.00	333	unidirectional a

7762155-7762487	0.00	333	divergent a/b

16 divergent; 15 unidirectional; 2 no annotated gene (33 total); div/uni= 1.07

**RNA9: CE11(C2), 1 month**

Region	p-value	Length	Locus type

6915-7268	0.00	354	divergent a/a

264245-264601	0.00	357	unidirectional a

782938-783288	0.00	351	unidirectional a

786356-786709	0.00	354	unidirectional a

1011139-1011492	0.00	354	unidirectional a

1037954-1038243	1.21E-6	290	unidirectional a

1039551-1039898	0.00	348	unidirectional a

1064683-1065007	8.19E-11	325	divergent a/b

1122014-1122364	0.00	351	divergent a/a

1208548-1208877	0.00	330	divergent a/b

1399269-1399601	0.00	333	divergent b/a

1402278-1402513	5.17E-18	236	divergent b/a

1403602-1403967	0.00	366	divergent b/a

1404548-1404874	0.00	327	divergent b/a

1600915-1601241	0.00	327	unidirectional a

1618161-1618580	1.15E-7	420	unidirectional a

2678070-2678405	0.00	336	divergent a/a

2882677-2882964	4.20E-10	288	unidirectional a

2897979-2898321	3.61E-8	343	unidirectional a

2918405-2918586	4.28E-9	182	divergent b/ves

3137303-3137650	0.00	348	no annotated gene

3396845-3397184	0.00	340	no annotated gene

5260614-5260991	0.00	378	divergent b/a

5456956-5457291	0.00	336	divergent b/a

5458074-5458335	0.00	262	divergent b/a

5458987-5459181	0.00	195	divergent b/a

6205332-6205667	0.00	336	unidirectional a

6216872-6217207	1.72E-8	336	divergent a/a

6218668-6219009	0.00	342	divergent a/a (pair)

7443575-7443907	0.00	333	unidirectional a

7762155-7762487	0.00	333	divergent a/b

12 divergent; 11 unidirectional; 2 no annotated gene (25 total); div/uni= 1.09

**RNA10: CE11(C2), 5 months**

Region	p-value	Length	Locus type

6915-7268	0.00	354	divergent a/a

243324-243551	0.00	228	unidirectional a

264245-264601	0.00	357	unidirectional a

285751-286083	0.00	333	divergent a/a

286759-287112	0.00	354	divergent a/a (a pair)

782938-783288	0.00	351	unidirectional a

785282-785626	0.00	345	unidirectional a

786356-786709	0.00	354	unidirectional a

1011139-1011492	0.00	354	unidirectional a

1037935-1038261	0.00	327	unidirectional a

1039551-1039898	0.00	348	unidirectional a

1055223-1055551	0.00	329	divergent b/a

1064702-1064889	2.81E-6	188	divergent a/b

1065065-1065243	0.00	179	divergent a/b

1122014-1122364	0.00	351	divergent a/a

1125759-1125858	5.81E-7	100	divergent a/a (a pair)

1133046-1133396	0.00	351	divergent b/a

1208548-1208877	0.00	330	divergent a/b

1399269-1399601	0.00	333	divergent b/a

1402184-1402513	6.20E-14	330	divergent b/a

1403602-1403967	0.00	366	divergent b/a

1404548-1404874	0.00	327	divergent b/a

1600915-1601241	0.00	327	unidirectional a

1618150-1618599	0.00	450	unidirectional a

2001903-2002139	5.05E-7	237	unidirectional a

2676785-2676969	0.00	185	divergent ves/ves

2678070-2678405	0.00	336	divergent a/a

2678837-2678963	2.69E-6	127	divergent ves/ves

2867318-2867671	0.00	354	divergent a/b

2897979-2898332	0.00	354	unidirectional a

2918405-2918586	0.00	182	divergent b/ves

3137303-3137650	0.00	348	no annotated gene

3396845-3397184	0.00	340	no annotated gene

5260614-5260991	0.00	378	divergent b/a

5262801-5263089	0.00	289	divergent b/a

5456956-5457291	0.00	336	divergent b/a

5458074-5458368	0.00	295	divergent b/a

5458922-5459254	0.00	333	divergent b/a

5467192-5467427	4.61E-14	236	divergent b/a

6205332-6205667	0.00	336	unidirectional a

6218668-6219009	0.00	342	divergent a/a

7421547-7421883	0.00	337	unidirectional a

7443575-7443907	0.00	333	unidirectional a

7762155-7762487	0.00	333	divergent a/b

20 divergent; 13 unidirectional; 2 with no annotated gene (35 total); div/uni= 1.54

**RNA11: CE11(C5), 1 month**

Region	p-value	Length	Locus type

6915-7268	0.00	354	divergent a/a

286759-287112	0.00	354	divergent a/a

782938-783288	0.00	351	unidirectional a

785282-785626	0.00	345	unidirectional a

786356-786709	0.00	354	unidirectional a

1011139-1011492	0.00	354	unidirectional a

1039551-1039898	0.00	348	unidirectional a

1055223-1055551	0.00	329	divergent b/a

1064683-1065243	0.00	561	divergent a/b

1122014-1122364	0.00	351	divergent a/a

1133046-1133396	0.00	351	divergent b/a

1399269-1399601	0.00	333	divergent b/a

1402277-1402513	2.05E-7	237	divergent b/a

1403602-1403967	0.00	366	divergent b/a

1404548-1404874	0.00	327	divergent b/a

1600915-1601241	0.00	327	unidirectional a

1618150-1618599	0.00	450	unidirectional a

2676785-2676969	0.00	185	divergent ves/ves

2678070-2678405	0.00	336	divergent a/a

2678755-2678964	2.35E-7	210	divergent ves/ves

2867318-2867671	0.00	354	divergent a/b

2882658-2882975	0.00	318	unidirectional a

2897990-2898213	1.76E-8	224	unidirectional a

2918405-2918586	2.29E-7	182	divergent b/ves

3137303-3137650	0.00	348	no annotated gene

3396845-3397184	0.00	340	no annotated gene

5260614-5260991	0.00	378	divergent b/a

5456956-5457291	0.00	336	divergent b/a

5458987-5459181	0.00	195	divergent b/a

6218668-6219009	0.00	342	divergent a/a

7421547-7421883	0.00	337	unidirectional a

7443575-7443907	0.00	333	unidirectional a

7762155-7762487	0.00	333	divergent a/b

16 divergent; 10 unidirectional; 2 no annotated gene (28 total); div/uni= 1.60

**RNA12: CE11(C5), 5 months**

Region	p-value	Length	Locus type

6915-7268	0.00	354	divergent a/a

285751-286083	0.00	333	divergent a/a

782938-783288	0.00	351	unidirectional a

785282-785626	0.00	345	unidirectional a

786356-786709	0.00	354	unidirectional a

1011139-1011492	0.00	354	unidirectional a

1037935-1038261	0.00	327	unidirectional a

1039384-1039539	7.17E-6	156	unidirectional a

1039551-1039898	0.00	348	unidirectional a

1055223-1055551	0.00	329	divergent b/a

1064683-1065243	0.00	561	divergent a/b

1122014-1122364	0.00	351	divergent a/a

1133046-1133396	0.00	351	divergent b/a

1208548-1208877	0.00	330	divergent a/b

1399269-1399601	0.00	333	divergent b/a

1402278-1402513	0.00	236	divergent b/a

1403602-1403967	0.00	366	divergent b/a

1404548-1404874	0.00	327	divergent b/a

1411516-1411704	0.00	189	unidirectional a

1600915-1601241	0.00	327	unidirectional a

1618150-1618599	0.00	450	unidirectional a

2001961-2002139	0.00	179	unidirectional a

2676785-2676969	0.00	185	divergent ves/ves

2678070-2678405	0.00	336	divergent a/a

2678770-2678964	2.51E-7	195	divergent ves/ves

2867318-2867671	0.00	354	divergent a/b

2897979-2898332	0.00	354	unidirectional a

2918405-2918586	0.00	182	divergent b/ves

3137303-3137650	0.00	348	no annotated gene

3396845-3397184	0.00	340	no annotated gene

5260614-5260991	0.00	378	divergent b/a

5456956-5457291	0.00	336	divergent b/a

5458987-5459181	0.00	195	divergent b/a

6205332-6205667	0.00	336	unidirectional a

6218668-6219009	0.00	342	divergent a/a

7421547-7421883	0.00	337	unidirectional a

7443575-7443907	0.00	333	unidirectional a

7761921-7762129	0.00	209	divergent a/b

7762155-7762487	0.00	333	divergent a/b

17 divergent; 13 unidirectional; 2 no annotated gene (32 total); div/uni= 1.31

^1^”p-value” indicates the statistical value for the alignment of the mapped sequence with that of the genomic locus to which it mapped.

^2^”Length” refers to the merged length of the mapped amplicons, in base pairs. This is shorter than the lengths of the primary amplicons because of having trimmed primer sequences from the ends.

^3^”Locus type” indicates whether the *ves* locus to which an amplicon mapped was comprised of a single *ves* gene (unidirectional), a *ves* gene in a divergent pair (divergent), or was one that was not annotated.

^4^”a”, “b”, “a/a”, “a/b”, and “b/a” refer to *ves* α, *ves* β, and pairs of *ves* genes in α/α or α/β pairs, and the order of the genes relative to genomic position. Underlining indicates the member of a gene pair to which an amplicon mapped.

Red, boxed sequences indicate *ves* loci which were assembled with multiple CKRD domains, and to which multiple amplicons mapped.
